# The Filterprep: A Simple and Efficient Approach for High-Yield, High-Purity Plasmid DNA Purification

**DOI:** 10.1101/2025.06.12.659418

**Authors:** Yu-Hsuan Cheng, Yu-Jiu Wu, Yung-Chun Shih, Yu-Qian Lin, Yu-Wei Chang, Yu-Heng Wu, En-Wei Hu, Ren-Hsuan Ku, Cheng-Mu Wu, Shao-Chi Wu, Ting-Yu Yeh, Chung-Te Chang

**Affiliations:** Institute of Biochemistry and Molecular Biology, National Yang Ming Chiao Tung University, Taipei 112, Taiwan; Department of Biotechnology and Laboratory Science in Medicine, National Yang Ming Chiao Tung University, Taipei 112, Taiwan

## Abstract

Plasmid DNA purification remains a fundamental yet resource-intensive step in molecular biology and biotechnology. Conventional protocols often yield plasmids with suboptimal purity, while commercial kits, though efficient, are costly and use proprietary formulations that limit transparency and customization. Here, we present Filterprep, a hybrid method combining classical ethanol precipitation with a single spin-column cleanup step, enabling rapid recovery of high-purity plasmid DNA with yields notably higher than those obtained using commercial kits, in approximately 40 minutes. Filterprep offers a transparent, cost-effective, and scalable alternative, simplifying workflows and reducing hands-on time without compromising DNA quality. The purified plasmids are compatible with a variety of downstream applications, including mammalian cell transfection, protein expression, and reporter assays. With its balance of efficiency and practicality, Filterprep is well suited for high-throughput laboratories and resource-limited settings.

## 1. Introduction

The isolation of plasmid DNA is a fundamental technique in molecular biology that has undergone significant evolution over the past several decades [1-4]. It primarily involves two phases: bacterial cell lysis and the selective recovery of plasmid DNA. While the initial cell disruption step, typically achieved through alkaline lysis, has remained relatively consistent, methods for plasmid DNA purification have diversified considerably [4].

Following cell lysis, traditional approaches relied heavily on alcohol-mediated DNA precipitation, followed by phenol-chloroform extraction to remove residual proteins and improve DNA purity [5-7]. Although these methods were cost-effective, they required substantial time and careful handling. Moreover, phenol and chloroform are hazardous organic solvents, making their routine use undesirable in modern laboratories. The advent of silica-based matrix technology introduced a transformative shift, enabling rapid processing through specialized spin columns [8]. These systems exploit the selective adsorption of DNA onto silica under specific chemical conditions, typically facilitated by ion-disrupting agents.

Modern commercial platforms now offer plasmid purification at various scales, from small-volume preparations to large-scale isolations. However, these systems involve inherent trade-offs among processing time, DNA yield, and economic cost. Plasmid recovery efficiency is influenced by multiple factors, including vector characteristics, bacterial host properties, and cultivation conditions. Moreover, commercial kits often employ proprietary buffer formulations, limiting researchers’ ability to troubleshoot and optimize protocols while increasing dependency on specific brands.

The continuous refinement of plasmid purification methodologies reflects the ongoing effort to balance procedural simplicity, time efficiency, and DNA yield. A notable advancement emerged with the Miraprep method, which demonstrated that ethanol incorporation prior to the silica-binding step could dramatically enhance DNA recovery [9]. This finding suggested that even well-established protocols retain untapped potential for further optimization.

Building on these insights, we developed a novel Filterprep method that strategically integrates elements of traditional precipitation techniques with contemporary column-based purification. Utilizing a single miniprep column yields substantial amounts of high-purity plasmid DNA while significantly reducing costs by eliminating the reliance on expensive commercial kits (**Table 1**). Filterprep effectively addresses the purity limitations of traditional methods and the high expenses associated with commercial platforms, providing researchers with a more accessible and efficient alternative. Thus, Filterprep offers a promising and user-friendly solution for high-quality, high-yield plasmid DNA isolation.

**Table 1.** Comparison of plasmid preparation methods using Qiagen kits and in-house protocols.

| Properties | Miniprep<br>(QIAGEN) | Midiprep<br>(QIAGEN) | Miraprep<br>(Pronobis et al., 2016) | Filterprep<br>(This Study) |
| --- | --- | --- | --- | --- |
| Bacteria culture (ml) | 1-5 | 25-100 | 50 | 50 |
| Max. column capacity (µg) | 20-25 | ~ 100 | N.A. | N.A. |
| Average yield (µg) | 23±5 | 176.7±21.75 | 531.6±49.79 | 1432±281.1 |
| Elution volume (µl) | 50-60 | variable | 150-175 | 200-400 |
| Time (min) | < 30 | 150 | < 30 | < 40 |
| Costs per µg DNA (cents) | 6.2 | 9.8 | 1.4 | 0.1 |
| N.A.: Not applicable as column capacity was no longer a limiting factor. |  |  |  |  |

## 2. Materials & Methods

### 2.1 Plasmids

The pEGFP-C1 plasmid (Clontech; 4.7 kb) encodes enhanced GFP. The FLuc plasmid (pCIneo- FLuc; 7.8 kb) encodes firefly luciferase.

### 2.2 Plasmid isolation kits

Filterprep was conducted using the spin column from the Miniprep kit of GeneAid (Presto™ Mini Plasmid Kit, Cat. No. PDH300), QIAGEN (QIAprep^®^ Miniprep Kit, Cat. No. 27106), Invitrogen by Thermo Fisher Scientific (PureLink™ Quick Plasmid Miniprep Kit, Cat. No. K210010), or Thermo Scientific (GeneJET Plasmid Miniprep Kit, Cat. No. K0502). The centrifuge spin filter columns contained a 0.22 µm filter mini column from NORGEN (298- 40000). The Midiprep kit from QIAGEN (Cat. No. 12145) was used for Midi preparations following the manufacturer’s instructions. Miraprep was performed using the QIAGEN Miniprep kit, following the previously described protocol [9].

### 2.3 Filterprep protocol

Detailed steps of the Filterprep are provided in the **Supplementary File 1**. Briefly, a 50 ml bacterial culture was inoculated with appropriate selective media and incubated on a shaker at 250 rpm at 37°C overnight. On the next day, the bacterial culture was transferred into a 50 ml tube and spun at 6000 xg at 4°C for 10 minutes. The supernatant was discarded, and the pellet was resuspended in 4 ml resuspension buffer with 100 mg/ml RNase I (QIAGEN) freshly added. 4 ml of lysis buffer was added to the bacterial suspension, and the tube was inverted 3-4 times. It was then incubated for 3 minutes at room temperature. 4 ml of neutralization buffer was added, and the tube was inverted 3-4 times. The bacterial lysate was spun at 13,200 xg at room temperature for 10 minutes. Next, filter the supernatants through a 20 ml syringe packed with a piece of Kimwipes™ into a 50 ml tube to remove residual debris remaining after centrifugation. The 1x volume of 95% ethanol (∼12 ml) was added to the supernatant and mixed thoroughly. The sample-ethanol mix was spun for 10 minutes at 13,200 xg.

After centrifugation, discard the supernatants and collect the pellets with 750 μl wash buffer. Transfer it into the spin column, spun at 13,200 xg at room temperature for 30 seconds, and the flow-through was discarded. These steps were repeated until the entire sample was passed through the spin columns. The column was washed two times with 750 μl wash buffer, spun after each wash at 13,200 xg at room temperature for 30 seconds, and the flow-through was discarded. The empty column was then spun one last time at 13,200 xg at room temperature for 1 minute to remove any residual wash buffer. After this, the old collection tube was discarded, and the column was put into a new tube. 200 μl of distilled water or elution buffer was added to the column, which was incubated for 2 minutes at room temperature, and spun at 13,200 xg for 2 minutes to elute the DNA from the column. After measuring the DNA concentration, the samples were stored at -20°C.

### 2.4. Cell culture

Human HEK293T cells were purchased from ATCC (CRL-3216) and were cultured in DMEM (Gibco 11995) supplemented with 10% (v/v) fetal bovine serum (Gibco) and grown at 37 °C with 5% CO2. The identity of these cells was authenticated through SRT profiling, and they were confirmed to be negative for Mycoplasma.

### 2.5. Immunofluorescence

For Immunofluorescence assays, HEK293T cells were seeded in 6-well dishes and transfected with plasmid DNA using Lipofectamine 2000 (Thermo). Cells were fixed and permeabilized as described previously, followed by immunostaining and quantification. [10, 11].

### 2.6. Western blot analysis

For Western blot assays, HEK293T cells were seeded in 10 cm dishes and transfected with 2.5, 5, or 10 μg pEGFP-C1 (Clontech) plasmid DNA using Lipofectamine 2000 (Thermo). The cells were washed 48 h after transfection with phosphate-buffered saline and lysed in 1 ml NET buffer (50 mM Tris-HCl (pH 7.5), 150 mM NaCl, 1 mM EDTA, 0.1% (v/v) Triton-X-100, 10% (v/v) glycerol). GFP and tubulin were detected using anti-GFP (Roche, 11814460001; 1:2,000) and anti-tubulin (Sigma, T6199; 1:5,000) antibodies, respectively. All Western blots were developed using the ECL Western blotting analysis system (GE Healthcare) as recommended by the manufacturer.

### 2.7. Luciferase assays

HEK293T cells were cultured in 6-well dishes and transfected with 2 μg of pCIneo-Fluc [12] construct using Lipofectamine 2000 (Thermo) according to the manufacturer’s protocol. 48 hours post-transfection, the cells were washed with phosphate-buffered saline and lysed in 1 ml NET buffer (50 mM Tris-HCl (pH 7.5), 150 mM NaCl, 1 mM EDTA, 0.1% (v/v) Triton-X-100, 10% (v/v) glycerol). Luciferase activity was measured using the Firefly assay buffer (100 mM Tris-HCl (pH 7.8), 15 mM MgCl_2_, 15 mM DTT, 0.6 mM Coenzyme A, 0.45 mM ATP, 4.2 mg/ml Luciferin). Firefly luciferase activity was normalized to total protein (A280) in cells transfected with the pCIneo-Fluc construct.

## 3. Results

Plasmid DNA isolation is a fundamental technique in molecular biology, with applications ranging from cloning to protein expression and gene therapy. Despite the widespread use of traditional and commercial plasmid isolation methods, significant limitations remain. Traditional methods are cost-effective and transparent, but often yield DNA of insufficient purity for downstream applications. In particular, protocols involving phenol–chloroform extraction pose significant safety hazards and are not ideal for routine or high-throughput use. In contrast, commercial kits provide high-purity DNA; however, they are expensive, rely on proprietary formulations, and limit scalability. These drawbacks highlight the need for a more accessible and efficient approach to plasmid isolation.

Among alternative methods, the Miraprep protocol combines a modified lysis procedure with spin-column purification, achieving high yields [9]. However, Miraprep requires multiple miniprep spin columns per preparation, increasing reagent consumption and cost. More critically, the mechanism underlying its high DNA recovery remains unclear. This uncertainty raises the question of whether high-efficiency DNA recovery could be achieved without proprietary components. A key insight into addressing this challenge comes from Pronobis et al., who demonstrated that a simple 0.22 μm centrifugal filter could effectively capture DNA when ethanol was added prior to filtration [9]. Their findings highlight the central role of ethanol precipitation in DNA recovery, shifting the focus away from silica-based binding.

Inspired by this insight, we sought to develop a plasmid isolation method that redefines the role of spin columns. Rather than relying on silica membranes for DNA binding, we hypothesized that spin columns could serve primarily as filtration tools, with ethanol precipitation driving DNA recovery. This approach aims to eliminate the need for costly chaotropic salts and proprietary reagents while maintaining high yields and purity.

### 3.1 The Filterprep: a novel approach to plasmid DNA isolation

To test this hypothesis, we developed the Filterprep method (**Figure 1**), a streamlined plasmid isolation protocol that combines conventional ethanol precipitation with selective purification using miniprep columns as filtration devices. By leveraging laboratory-prepared buffers with fully transparent compositions, Filterprep provides a cost-effective and efficient alternative to commercial kits. The key innovation of Filterprep lies in its strategic use of miniprep columns: rather than relying on complex chemical interactions, we treat the columns as precise filtration systems to further purify ethanol-precipitated DNA. Notably, while Miraprep requires five miniprep columns for optimal DNA recovery, Filterprep achieves comparable results using only a single column, significantly reducing both material costs and procedural complexity. This approach not only simplifies the plasmid isolation process but also maintains high yields and purity, making it accessible to researchers in resource-limited settings.

**Figure 1.**
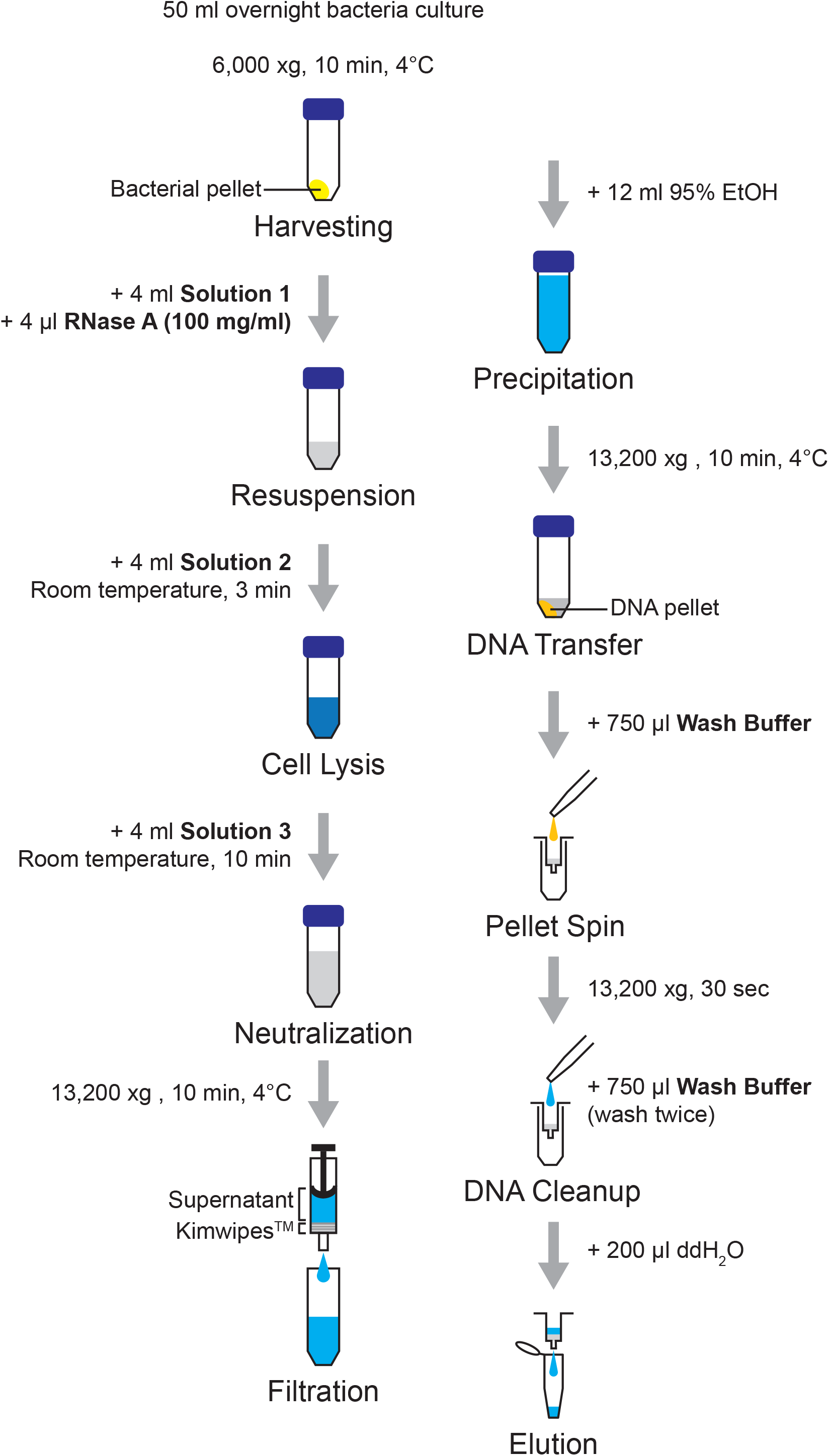
Schematic diagram of the Filterprep method. Overview of the plasmid DNA purification workflow showing bacterial cell harvesting, lysis, debris clearance, ethanol precipitation, column-based purification, and final elution steps. Critical parameters, including centrifugation speeds, incubation times, and buffer volumes, are specified for each step.

To validate our approach, we performed a comprehensive comparison of Filterprep with commercial Midiprep kits and the Miraprep method (**Figure 2A**). Our analysis demonstrated that Filterprep consistently yielded approximately 1432±281 μg of plasmid DNA, exceeding the output of commercial kits while maintaining high purity. Spectrophotometric analysis showed A260/A280 ratios between 1.8 and 2.0, comparable to those obtained with standard commercial preparations. Furthermore, gel electrophoresis confirmed the integrity and purity of the isolated DNA, revealing a banding pattern closely resembling that of DNA purified using commercial kits (**Figure 2B**).

**Figure 2.**
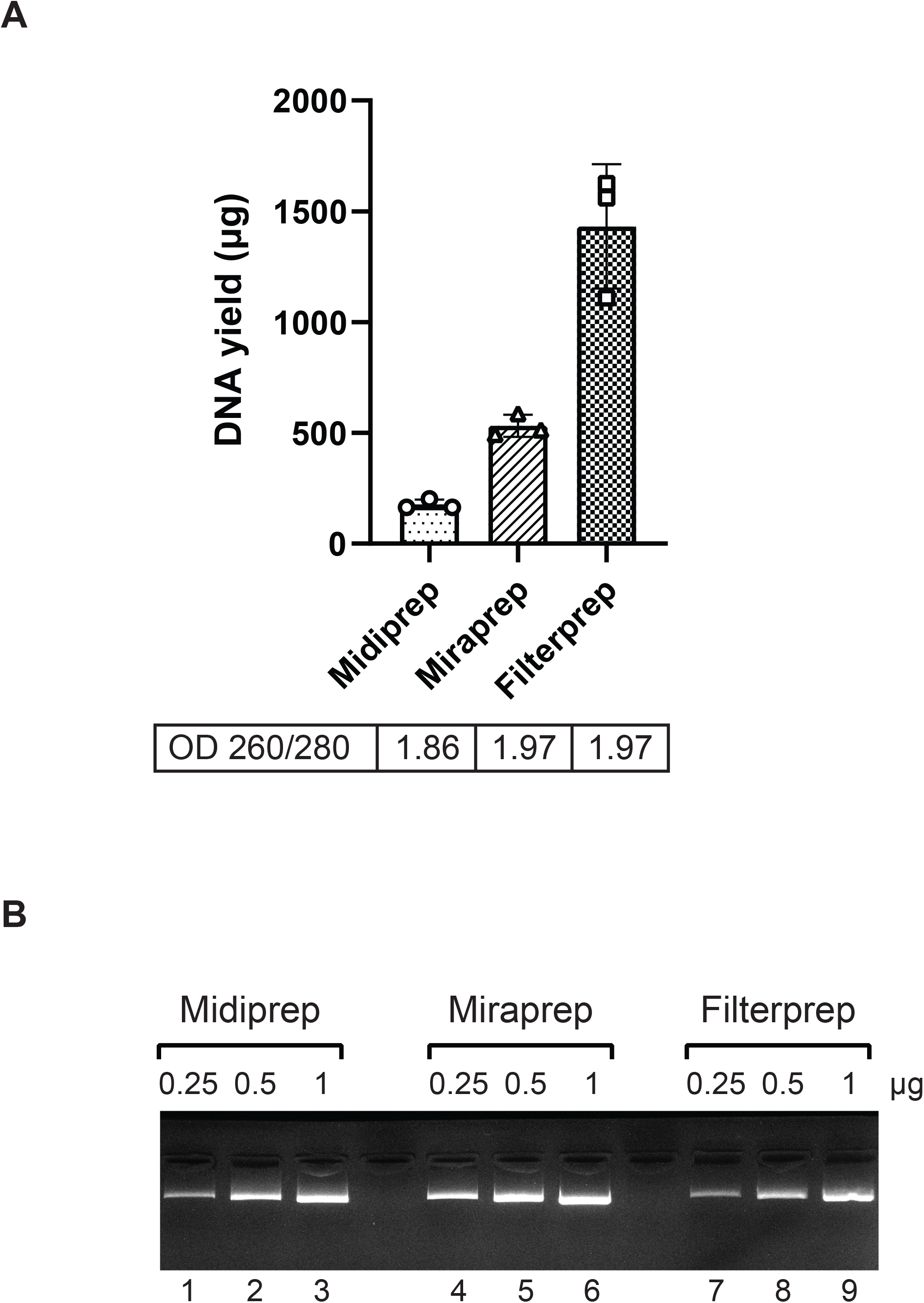
Superior yield and purity performance of Filterprep compared to commercial kits and Miraprep. (A) Yield and quality of DNA from Midiprep, Miraprep, and Filterprep. DNA yields are presented as mean values with standard deviation calculated from three independent experiments and average OD260/280 at the bottom. (B) Agarose gel electrophoresis (1%) of the plasmid DNA. Samples from different methods (Lane 1-3: Midiprep; Lane 4-6: Miraprep; Lane 7-9: Filterprep) with increasing concentration (0.25, 0.5, 1 µg) were electrophoresed on agarose gel and visualized by ethidium bromide staining.

Critically, these improved results were achieved using only standard laboratory materials, elevating our approach from a simple technical adjustment to a practical and impactful alternative. By providing an accessible and cost-effective substitute for commercial plasmid isolation kits, Filterprep offers particular value in resource-limited settings. Overall, our redesign streamlines DNA recovery while preserving high standards of purity and yield. This work underscores how re-evaluating established protocols can lead to meaningful innovations through thoughtful simplification and optimization.

### 3.2 Performance of Filterprep with Different Brands and Types of Columns

To evaluate the robustness and versatility of the Filterprep method, we examined whether the choice of filtration tools, including various brands of miniprep columns and alternative systems, would affect plasmid DNA yield and quality. We tested four commercially available miniprep columns (Geneaid, QIAGEN, Invitrogen, and Thermo), a 0.22 µm non-silica membrane filter column (NORGEN), and used a commercial midiprep kit (QIAGEN) as a reference.

Our analysis showed no significant differences in DNA yield or purity among the different miniprep columns or the 0.22 µm filter column (**Figure 3A**). All approaches consistently yielded approximately 1316±173 µg of plasmid DNA, with OD260/280 ratios ranging from 1.8 to 2.0, comparable to those obtained using the midiprep kit. These findings indicate that the performance of Filterprep does not depend on the brand of column or the use of silica membranes but instead relies on the efficiency of the preceding ethanol precipitation step.

**Figure 3.**
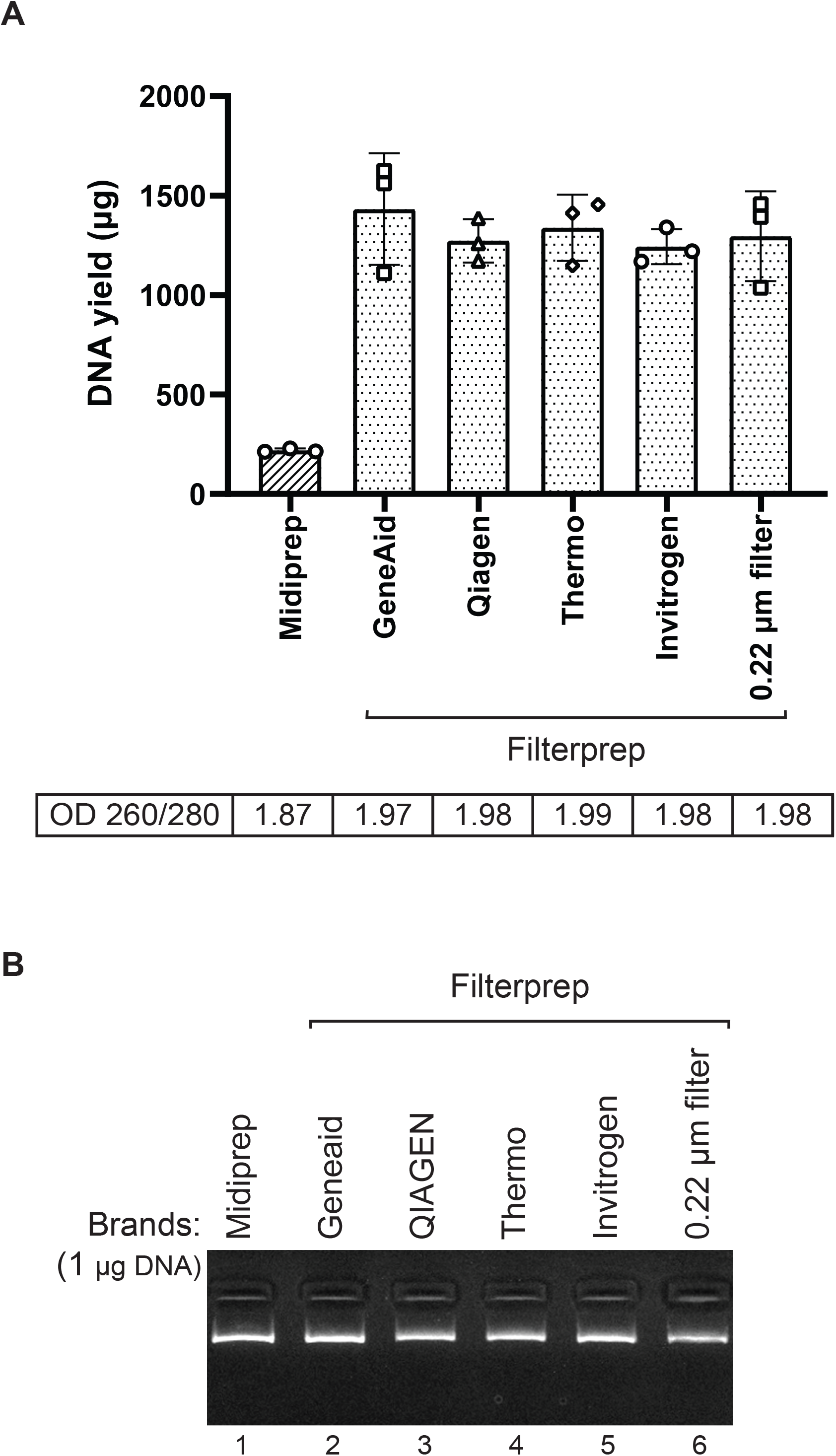
Filterprep performance across different brands and types of columns. (A) Yield and quality of DNA from four commercially available miniprep columns or the 0.22 µm filter column. DNA yields are presented as mean values with standard deviation calculated from three independent experiments and average OD260/280 at the bottom. (B) Agarose gel electrophoresis (1%) of the plasmid DNA. Samples from different columns with specific concentration (1 µg) were electrophoresed on agarose gel and visualized by ethidium bromide staining.

To confirm these results, we performed agarose gel electrophoresis to assess DNA integrity and quality (**Figure 3B**). The gel profiles showed consistent banding patterns across all tested methods, with clear signals corresponding to both supercoiled and open-circular plasmid forms. No quality differences were observed among the miniprep columns, the 0.22 µm filter column, and the midiprep control, reinforcing the robustness of the Filterprep protocol.

These results emphasize the flexibility and cost-effectiveness of the Filterprep method. The ability to use alternative filtration systems without compromising DNA yield or quality makes it particularly suitable for laboratories with limited access to specialized columns. Furthermore, the consistent performance across different column brands supports the reliability of Filterprep as a broadly applicable and practical alternative to commercial plasmid purification kits.

### 3.3 Functional Validation of Filterprep-Isolated Plasmid DNA

To demonstrate the purity and functionality of plasmid DNA isolated using the Filterprep method, we performed DNA sequencing and mammalian cell transfection experiments, comparing the results to those obtained with plasmid DNA purified using a commercial midiprep kit.

We first evaluated the quality of Filterprep-isolated plasmid DNA by performing Sanger sequencing. As shown in **Figure 4A**, the sequencing chromatograms of plasmids purified using Filterprep exhibited clear, high-intensity peaks with minimal background noise, comparable to those obtained from plasmids isolated with the commercial midiprep kit. The high-quality sequencing results confirm that Filterprep-isolated plasmids are free of contaminants that could interfere with downstream applications, such as residual salts or proteins, and are suitable for sequencing-based analyses.

**Figure 4.**
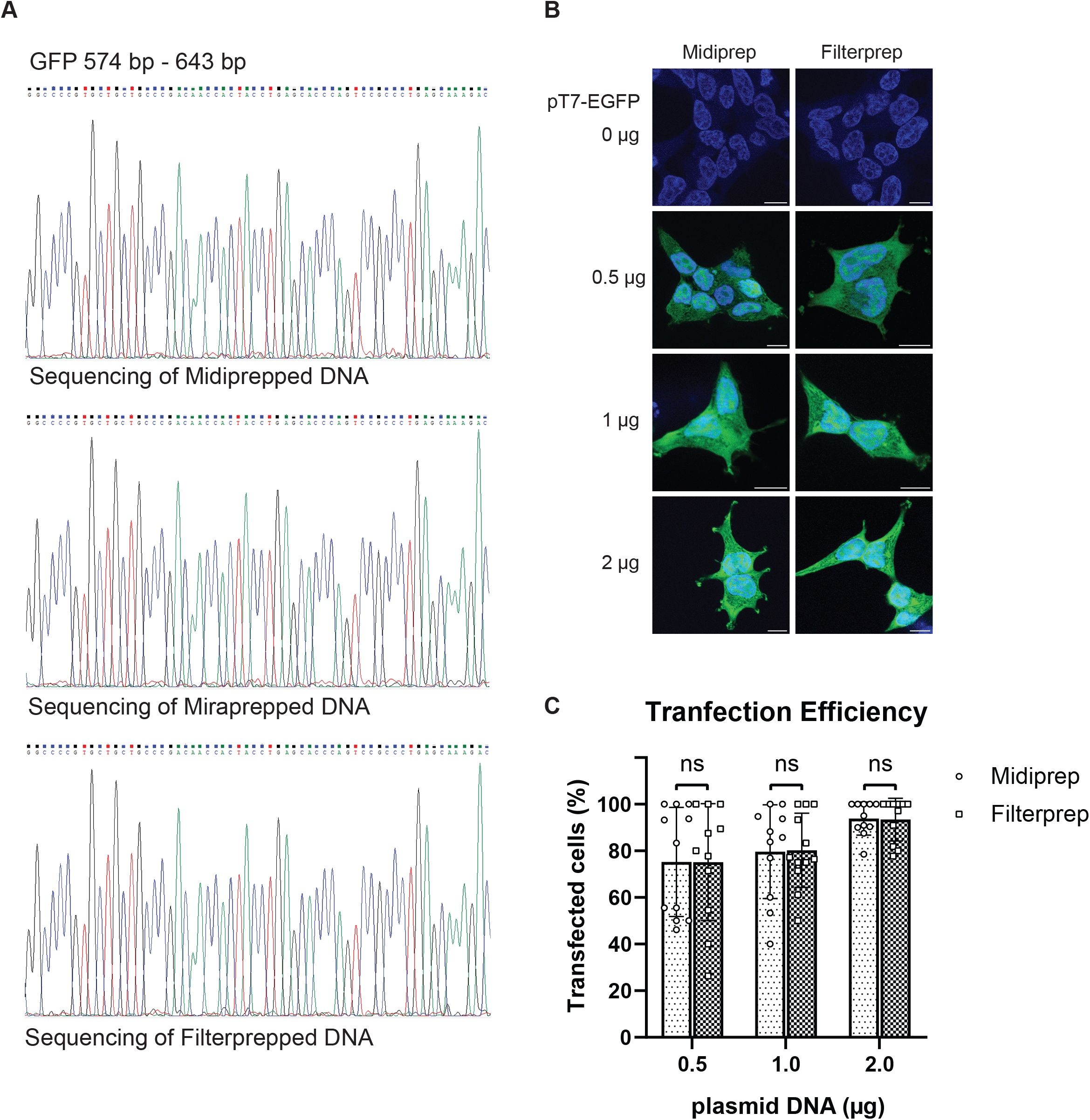
Filterprep-isolated plasmids can be used for sequencing and transfection. (A) Sequencing reaction of Midiprepped, Miraprepped, and Filterprepped GFP plasmid. The sequence from 574 base pairs (bp) to 643 bp is shown. 1 µg of DNA was used for the sequence reaction. (B) HEK293T wild-type cells overexpressing GFP (green) were fixed, then counterstained with DAPI to visualize the nucleus (blue). Scale bar, 10 μm. (C) Quantification of transfection efficiency in the HEK293T cells overexpressing GFP. Ten fields of view for each cell line (>100 cells) were used to quantify the number of transfected cells. Data represents the mean values with standard deviation, and the significance was evaluated with two-way ANOVA.

Next, we assessed the functionality of Filterprep-isolated plasmid DNA in mammalian cell transfection experiments. We transfected HEK293T cells with 0, 0.5, 1, and 2 µg of a GFP-expressing plasmid purified using either Filterprep or the commercial midiprep kit. At 48 hours post-transfection, we visualized GFP expression using fluorescence microscopy (**Figure 4B**). Both Filterprep and midiprep-isolated plasmids produced robust and dose-dependent GFP expression, with no observable differences in fluorescence intensity between the two methods.

To quantify the transfection efficiency, we performed fluorescence intensity measurements of the transfected cells (**Figure 4C**). The results revealed no significant differences in GFP expression levels between cells transfected with Filterprep-isolated plasmids and those transfected with midiprep-isolated plasmids at all tested concentrations. This indicates that Filterprep-isolated plasmids are equally effective in driving transgene expression in mammalian cells, further validating their suitability for functional assays.

Together, these experiments demonstrate that plasmid DNA isolated using the Filterprep method is of high quality and fully functional for standard molecular and cell biology applications. The sequencing results confirm the absence of contaminants that could interfere with downstream analyses, while the transfection experiments highlight the ability of Filterprep- isolated plasmids to efficiently drive gene expression in mammalian cells. These findings solidify Filterprep as a reliable and cost-effective alternative to commercial plasmid isolation kits, particularly for researchers requiring high-quality DNA for functional studies.

### 3.4. Versatility of Filterprep-Isolated Plasmid DNA in Routine Biochemical Applications

To further validate the utility of Filterprep-isolated plasmid DNA in routine biochemical experiments, we conducted a series of functional assays, including Western blot analysis, isolation of plasmids of varying sizes, and luciferase activity assays. These experiments were designed to demonstrate the broad applicability of Filterprep in standard laboratory workflows.

We first evaluated the expression efficiency of proteins encoded by plasmids isolated using Filterprep. HEK293T cells were transfected with the EGFP plasmid purified using either Filterprep or a commercial midiprep kit. At 48 hours post-transfection, protein expression was analyzed by Western blot (**Figure 5A**). The results revealed comparable levels of EGFP expression between cells transfected with Filterprep-isolated plasmids and those transfected with midiprep-isolated plasmids. This confirms that Filterprep-isolated plasmids are fully capable of driving efficient protein expression, making them suitable for downstream protein analysis applications.

**Figure 5.**
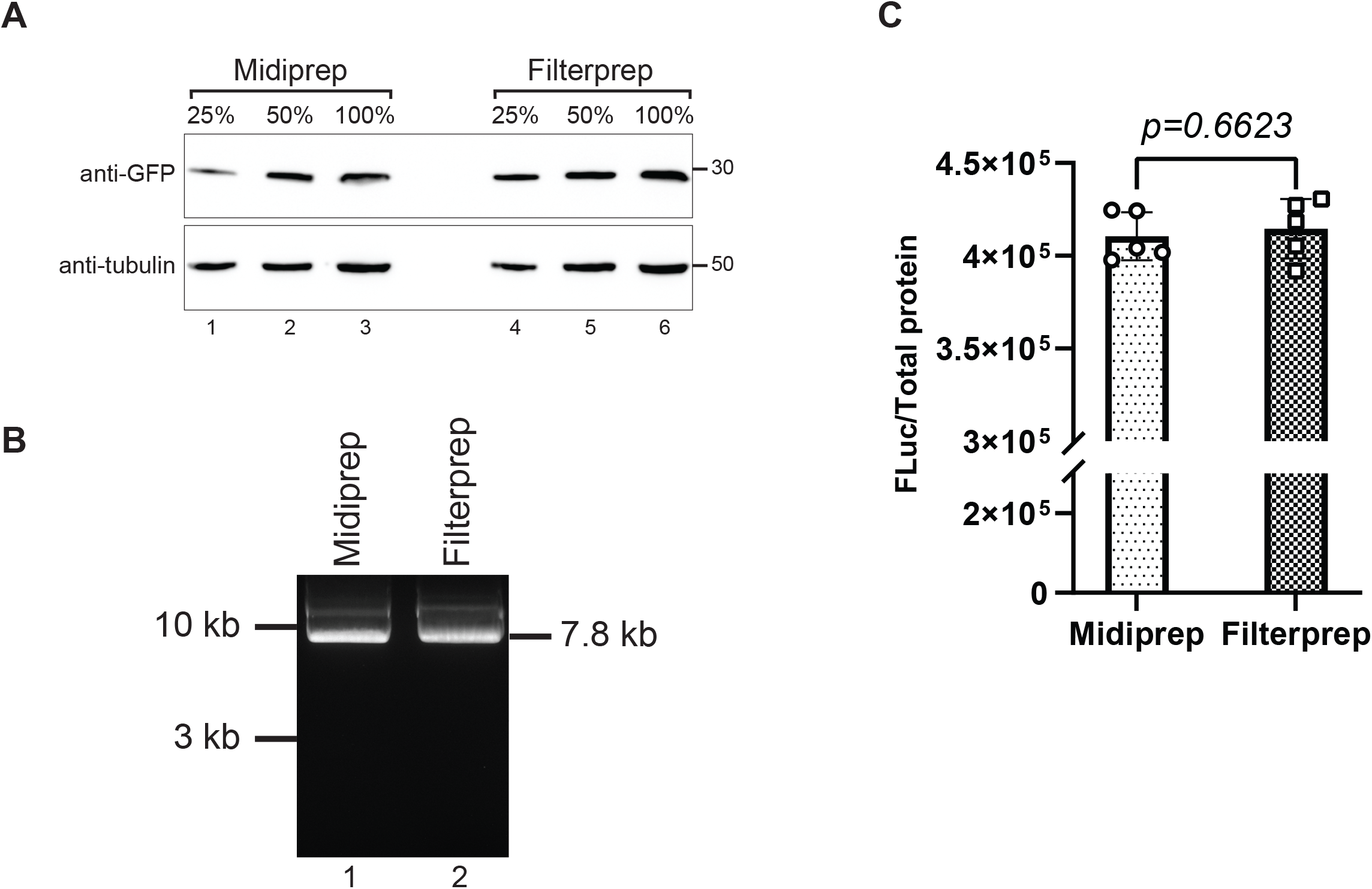
Filterprep-isolated plasmids are functional in driving reporter gene expression. (A) Midiprepped and Filterprepped GFP protein levels in HEK293T wild-type cells were examined by immunoblotting using antibodies against GFP and tubulin, the latter serving as a loading control. (B) Agarose gel electrophoresis (1%) of the plasmid DNA. Samples from different methods with specific concentration (1 µg) were electrophoresed on agarose gel and visualized by ethidium bromide staining. (C) Luciferase activity assessment by luciferase reporter assay in HEK293T cells overexpressing Midiprep or Filterprep-isolated plasmids. Total protein (A280) was used as an internal control. Data represents mean values with standard deviation from five independent experiments. Statistical significance was determined using an unpaired t-test.

Having previously used the EGFP plasmid (pEGFP-C1; 4.8 kb) to validate the effectiveness of Filterprep, we next assessed its performance on a larger construct. The FLuc plasmid (pCIneo- FLuc; 7.8 kb) was processed using both Filterprep and a commercial midiprep kit, and the resulting DNA was examined by agarose gel electrophoresis. As shown in **Figure 5B**, both preparations exhibited clear and consistent banding patterns, indicating that Filterprep can effectively isolate high-quality plasmid DNA across different plasmid sizes. These results support its applicability for a broad range of plasmid constructs.

To evaluate the functional integrity of Filterprep-isolated plasmids, we performed luciferase activity assays using HEK293T cells transfected with Fluc plasmids isolated by either Filterprep or the commercial midiprep kit. Luciferase activity was measured at 48 hours post-transfection (**Figure 5C**). The results showed that cells transfected with Filterprep-isolated plasmids exhibited luciferase activity levels comparable to, and in some cases slightly higher than, those transfected with midiprep-isolated plasmids. This indicates that Filterprep-isolated plasmids are not only highly pure but also fully functional in driving reporter gene expression.

Together, these experiments demonstrate that Filterprep-isolated plasmid DNA is highly versatile and suitable for a wide range of routine biochemical applications. The Western blot analysis confirms its ability to support efficient protein expression, while the isolation of plasmids of different sizes highlights its robustness across various experimental needs. Additionally, the luciferase activity assays underscore the functional integrity of Filterprep- isolated plasmids, making them a reliable and cost-effective alternative to commercial kits for diverse molecular and biochemical studies.

## 4. Discussion

The Filterprep method represents a transformative approach to plasmid DNA isolation, challenging conventional paradigms in molecular biology laboratory techniques. By reimagining the roles of miniprep columns and ethanol precipitation, we demonstrate that high-quality DNA purification can be achieved through a simpler, more economical strategy (**Table 1**).

Currently, most research laboratories rely on commercial kits for plasmid DNA isolation. Although effective, these kits utilize complex chemical interactions, such as chaotropic salts, to facilitate DNA binding and purification. However, their high costs can burden laboratories, especially those conducting high-throughput experiments or operating under limited funding. Furthermore, the proprietary nature of commercial formulations conceals the exact composition of buffers and reagents, complicating troubleshooting, optimization, and customization for specific experimental needs. Filterprep overcomes these challenges by applying fundamental molecular separation principles while maintaining or even enhancing DNA purity and yield.

A key innovation of Filterprep lies in its strategic redefinition of the DNA isolation workflow. By recognizing that miniprep columns can function primarily as filtration tools rather than platforms for complex chemical interactions, we have simplified the process without compromising performance. This approach not only reduces costs but also increases accessibility, making high-quality plasmid isolation feasible for a wider range of researchers. Importantly, Filterprep relies on laboratory-prepared buffers with fully transparent compositions, eliminating the dependency on proprietary reagents and providing researchers with complete control over the isolation process.

The robustness of Filterprep is underscored by its consistent performance across a variety of experimental validations. Our results demonstrate that plasmids purified using Filterprep are suitable for a wide range of downstream applications, including DNA sequencing, immunofluorescence, Western blotting, and luciferase assays. High-quality DNA preparations, characterized by optimal spectrophotometric ratios and significant yields, were consistently achieved and found to be comparable to those obtained using commercial kits.

Economic considerations further highlight the advantages of Filterprep. By significantly reducing reagent costs and eliminating the need for expensive commercial kits, this method democratizes access to high-quality plasmid DNA isolation. This is particularly impactful for research institutions in resource-limited settings, where financial constraints often hinder the adoption of advanced molecular biology techniques.

Looking ahead, several avenues for further optimization and validation of Filterprep can be explored. Future studies could focus on refining precipitation conditions, investigating alternative filtration approaches, or adapting the method for specific plasmid types or bacterial strains. Additionally, comparative studies examining long-term DNA stability and performance under various storage conditions would provide further validation of the method’s robustness and reliability.

Given its simplicity, scalability, and ability to yield high-quality plasmid DNA, the Filterprep method could serve as a valuable platform in applications beyond basic research. In particular, large-scale plasmid preparations are increasingly required in DNA vaccine development, viral vector production, and synthetic biology workflows where cost-effective and high-throughput plasmid purification is critical. The demonstrated compatibility with downstream applications such as transfection, protein expression, and reporter assays suggest that Filterprep may be especially useful in educational labs, resource-limited settings, and industrial pipelines where consistent plasmid DNA quality and yield are essential.

In conclusion, Filterprep offers a practical, cost-effective, and efficient alternative to traditional plasmid isolation methods. By simplifying the workflow, reducing dependency on proprietary reagents, and providing full transparency in buffer composition, this method has the potential to significantly impact molecular biology research, particularly in resource-constrained settings.

## ACKNOWLEDGMENTS

We are grateful to Jing-Yi Siao for her contributions to project administration and investigation during the early stages of the project.

